# Sibling rivalry: Males with more brothers develop larger testes

**DOI:** 10.1101/332775

**Authors:** Heidi S. Fisher, Kristin A. Hook, W. David Weber, Hopi E. Hoekstra

**Author notes:** Corresponding author: Heidi S. Fisher.

## Abstract

When females mate with multiple partners in a reproductive cycle, the relative number of competing sperm from rival males is often the most critical factor in determining paternity. Gamete production is directly related to testis size in most species, and is associated with both mating behavior within a system and perceived risk of competition. *Peromyscus maniculatus* is naturally promiscuous and males invest significantly more in sperm production than males of *P. polionotus*, their monogamous sister-species. Here we show that the relatively larger testes in *P. maniculatus* are retained, even after decades of enforced monogamy in captivity. While these results suggest that differences in sperm production between species with divergent evolutionary histories can be maintained, we also show that the early rearing environment of males can strongly influence their testis size as adults. Using a second-generation hybrid population to increase variation in testis size, we show that males reared in litters with more brothers develop larger testes as adults. Importantly, this difference in testis size is also associated with increased fertility. Together, our findings suggest that sperm production may be both broadly shaped by natural selection over evolutionary timescales and also finely tuned during early development.

## Introduction

In systems in which females mate multiply, the competition among sperm from different males to fertilize the available ova can be intense and drive the evolution of male reproductive traits (Parker 1970). Sperm competition theory predicts that males that can mate more frequently, produce larger, more competitive ejaculates and/or a greater number of sperm will achieve greater reproductive success (Parker and Pizzari 2010). Sperm production, however, is energetically costly (Dewsbury 1982) and can come at the expense of other important features including the onset of reproductive maturity, immune function, survival, and secondary sexual traits (e.g., Boonstra et al. 2001; Tuni et al. 2016; Simmons et al. 2017). Therefore, while increased investment in sperm production is advantageous in a competitive context, males can benefit from prudent reproductive allocation when competition is relaxed (e.g., Pitnick et al. 2006; Ramm and Stockley 2007; 2009; Firman et al. 2013). Populations that have evolved under divergent mating systems, and consequently vary in levels of sperm competition, provide insight into how social conditions affect an individual’s reproductive investment and, ultimately, fitness (Parker 2002; Firman and Simmons 2011).

*Peromyscus* rodents exhibit striking variation in testicular, ejaculate and sperm traits across species (Linzey and Layne 1969; 1974), which are associated with differences in mating strategy (Turner et al. 2010; Bedford and Hoekstra 2015). This trend is perhaps best exemplified by two sister-species, the deer mouse (*P. maniculatus*) and the oldfield mouse (*P. polionotus*). In *P. maniculatus*, both sexes mate with multiple partners, often in overlapping series just minutes apart (Dewsbury 1985), and females frequently carry multiple-paternity litters in the wild (Birdsall and Nash 1973); by contrast, *P. polionotus* is strictly monogamous, as established from both behavioural (Dewsbury 1981) and genetic data (Foltz 1981). Consistent with the well-documented relationship between relative testis size and level of sperm competition in rodents (Ramm et al. 2005), wild-caught *P. maniculatus* have testes roughly twice as large as those observed in *P. polionotus* males (Linzey and Layne 1969). Testis size has been shown to increase with population density (Long and Montgomerie 2006) and longer breeding seasons (Ribble and Millar 1992) in *P. maniculatus*, but it is less clear if species-specific differences are retained through multiple generations of enforced monogamy in captivity. Moreover, while there is evidence in other rodents that gamete production can be phenotypically plastic in response to sperm competition risk (Ramm and Stockley 2009; Firman et al. 2013; Klemme et al. 2014), it remains unclear how early in development males can respond to social cues either by a directed response or by passive uptake of circulating hormones.

Here we show that despite over six decades of enforced monogamy, male *P. maniculatus* have retained the reproductive morphology that enables them to produce more sperm than their naturally monogamous congener, *P. polionotus*. Then, using a second-generation hybrid intercross of these *Peromyscus* species to generate a greater range of reproductive phenotypes (Fisher et al. 2016; Bendesky et al. 2017) and to breakup co-adapted gene complexes (West-Eberhard 2003), we examine whether the social composition of the early developmental environment affects sperm production in adults and impacts reproductive potential.

## Materials and methods

### EXPERIMENTAL ANIMALS

*Peromyscus maniculatus bairdii* and *P. polionotus subgriseus* were originally obtained from the Peromyscus Genetic Stock Center at the University of South Carolina and then maintained at Harvard University and the University of Maryland. Founders of the *P. maniculatus* colony were collected in 1946 and 1948, and *P. polionotus* in 1952 (Bedford and Hoekstra 2015). Since establishment, the colonies have been outbred and maintained via monogamous pairings; non-breeders housed with same-sex conspecifics (Foster 1959). All mice in this study were maintained at 22°C and a 16:8 LD cycle to control for effects of photoperiod on testicular development and onset of puberty (Whitsett et al. 1984).

### REPRODUCTIVE MEASURES

We used sexually-mature *P. maniculatus* (*N*=58) and *P. polionotus* (*N*=35) males to examine reproductive morphology of the captive stocks. After sacrifice via isoflurane overdose, we weighed the male, immediately removed and weighed each testis, and then photographed each testis on 1cm grid paper. We estimated the two-dimensional area from the images using ImageJ 1.x (Schindelin et al. 2015) and the “freehand” drawling tool. We calculated the mean testis mass and area for each male. To account for body size differences between males within these two species (Table 1), we included body mass as a covariate in our analyses, a method better suited to estimating relative testis size rather than using the ratio of testis to body mass or residuals (García Berthou 2001; Lupold et al. 2009).

**Table 1.** Body and testis size of focal species.

| Species | Body mass (g)<br>mean ± SE | Testis mass (mg)<br>mean ± SE | Testis area (mm<sup>2</sup>)<br>mean ± SE |
| --- | --- | --- | --- |
| *Peromyscus maniculatus* | 21.0 ± 0.32 | 123.2 ± 3.9 | 36.7 ± 1.1 |
| *Peromyscus polionotus* | 14.4 ± 0.40 | 70.1 ± 3.6 | 24.6 ± 0.9 |

To produce the hybrid population to examine the effects of early developmental environment on reproductive investment, we intercrossed two *P. polionotus* males and two *P. maniculatus* females to produce 38 first-generation hybrids, and then mated siblings to generate 300 second-generation (F_2_) hybrid male progeny. To test for an effect of testis size on fertility, we weaned each of the F_2_ hybrid males from their parents at 25 days of age and housed them with same-sex littermates until they were sexually mature and at least 60 days old. We then housed each male with a F_2_ female chosen at random from the same grandparents as the male for at least 7 days. Being paired with a female reduced male phenotypic variability that might arise from dominance hierarchies among male littermates and minimized differences in reproductive condition among males by exposing all of them to a female in estrus (*Peromyscus* estrous cycle is 5 days [Asdell 1964]); these females were monitored for at least 30 days after being paired with a male to determine his ability to fertilize a female (i.e., whether he sired offspring or not); note that this is conservative estimate which does not account for variation in female fertility nor missed litters due to offspring mortality and cannibalism. Following methods above, we measured testis area of each F_2_ male, which correlates tightly with testis mass (see Results) as a measure of testis size. We found no effect of male age (range: 71-183 days) or pairing duration (range: 7-108 days) on male fertility nor testis size (see Results).

### STATISTICAL ANALYSES

All statistical analyses were performed using R version 3.4.2 (R Core Team 2018). We used linear models (LM) to compare testis size between the two focal species, and to determine whether pairing duration with a female or male age differed between F_2_ sires and non-sires. To investigate the effects of social condition on testis size of males, we used a linear mixed model (LMM) using the lmer function from the “lme4” R package (Bates et al. 2014). The only random effect that contributed to variation in the response variable (i.e., mean testis area) was the grandparental line from which the hybrids were derived. This random factor was then used in bivariate analyses for predictors of interest, including: number of male littermates, number of female littermates, proportion of males in litter, litter order, litter size, age, time paired with female, mate identification, and fertility. Predictors that had a *p≤*0.20 were considered for the final model and were screened for collinearity using a linear model. Variables that were collinear with other variables of greater relative significance were removed from the final model, which resulted in only two remaining predictors: the number of male littermates and fertility.

## Results

We found that captive *P. maniculatus* males have significantly greater testis mass (LM: F_2,90_=61.28, *p*<0.001; Table 1) and testis area (LM: F_2,89_=38.56, *p*<0.001; Figure 1, Table 1) than *P. polionotus* males. Moreover, testis mass and area are directly correlated in both *P. maniculaus* (LM: F_1,55_=38.0, *p*<0.001) and *P. polionotus* (LM: F_1,33_=7.76, *p*=0.008).

**Figure 1.**
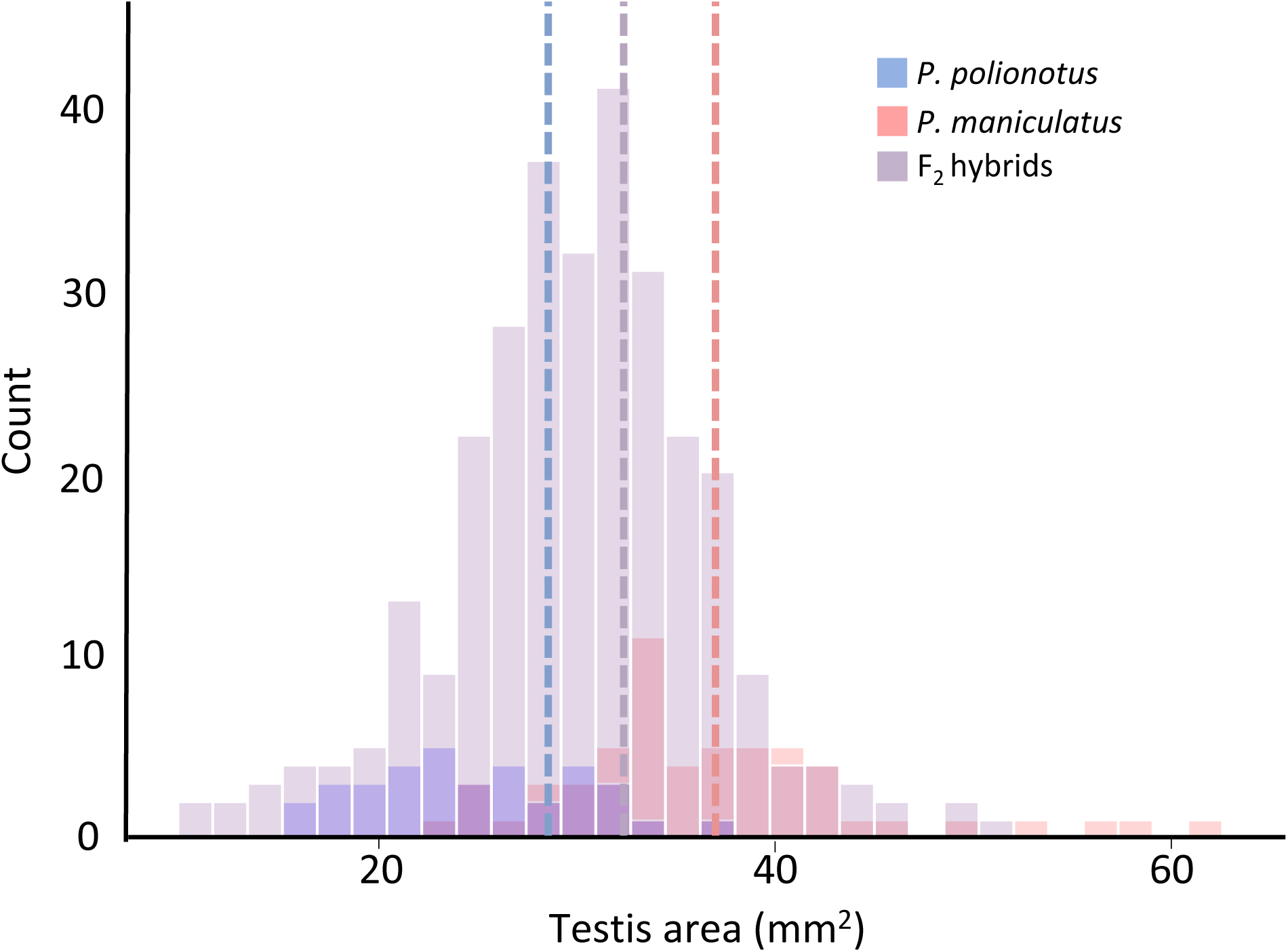
Frequency distribution of male mean testis size of the three study populations: *P. maniculatus* (*N*=58), *P. polionotus* (*N*=35), and F_2_ hybrids (*N*=300). The population mean testis size of each population is demarcated by dotted lines. Note truncated x-axis.

We found a significant effect of the social composition during development on the adult testis size in *P. maniculatus* x *P. polionotus* F_2_ hybrid males. Males reared with more brothers have significantly larger testes than those reared with fewer brothers (LMM: *N=*300, *p*<0.001; Figure 2a, Table 2). Additionally, males who successfully produced offspring after being paired with a female have significantly larger testes than males that did not sire offspring, indicating a significant effect of testis size on fitness (LMM: *N=*300, *p=0.024*; Figure 2b, Table 2). We found no effect of male age on male fertility (LM: *N=*85, *p=*0.69) or on testis size (LMM: *N=300, p=*0.70). Similarly, we found no effect of pairing duration on fertility (LM: *N=*85, *p=*0.34) or testis area (LMM: *N=300, p=*0.47).

**Table 2.** Body and testis size of focal species.

|  | Testis area (mm<sup>2</sup>) |  |  |
| --- | --- | --- | --- |
| | $N=300$ | | |
| Model term | Beta ± SE | *t* | *p* |
| Intercept | 26.97 ± 0.77 | | |
| Number of male littermates | 1.49 ± 0.31 | 4.81 | 2.40e-06 |
| Fertility | 1.84 ± 0.81 | 2.27 | 0.0237 |

**Figure 2.**
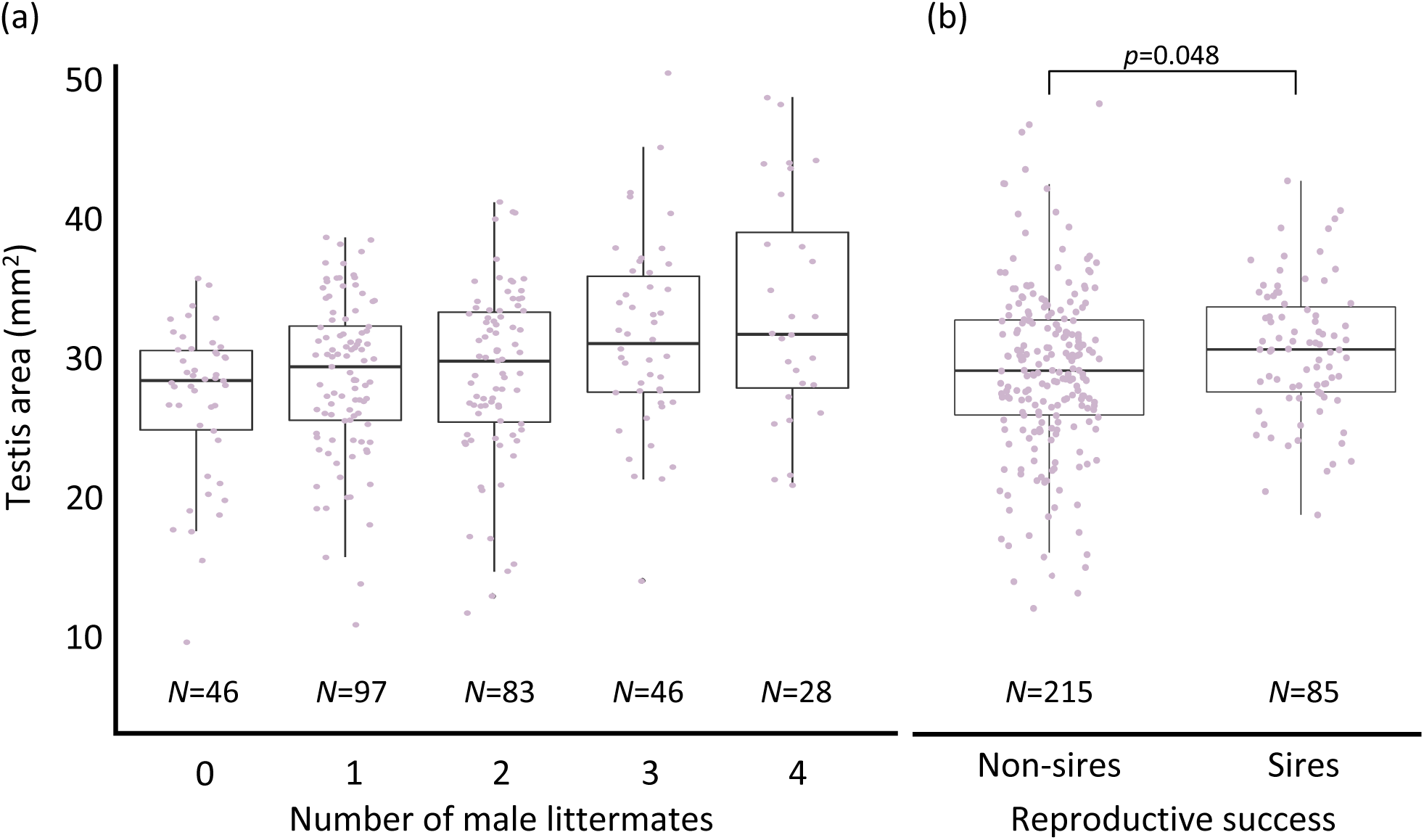
Male mean testis size of F_2_ hybrids. (a) Males with more male litter mates have significantly larger testes than those reared with fewer brothers (LMM: *N=*300, *p*<0.001). (b) F_2_ hybrid males that sired offspring after being paired with a female have larger testes than males that did not sire. Box-plots represent median and interquartile ranges with raw data (male mean testis size) overlaid in purple. Sample sizes are provided below data. Note truncated y-axis.

## Discussion

Our results suggest that both evolutionary history and social cues during development can have an important effect on investment in sperm production in *Peromyscus* mice. First, we confirmed that males of the naturally promiscuous species, *P. maniculatus* invest significantly more in sperm production compared to their monogamous sister-species, *P. polionotus,* despite decades of captive rearing under enforced monogamy. Although the relative difference in testis size is not as profound as observed in wild-caught males (Linzey and Layne 1969), the trend remains and suggests there are interspecific genetic differences that modulate reproductive development. Second, we found that the social environment can also influence investment in sperm production. Males exposed to more brothers early in development have larger testes and sire offspring as adults. Finally, our data show that testis size is significantly and positively associated with sperm production in the focal species, and ultimately with fertility.

One potential explanation for why we find that males with more brothers are likely to have larger testes is that there may be a maternal effect on both offspring “quality” and sex ratio; for example, females in greater condition may be able to produce more competitive sons and may therefore selectively reabsorb or cannibalize litters to bias the sex ratio in favor of males. The Trivers-Willard hypothesis (Trivers and Willard 1973) predicts that systems in which males have higher reproductive variance than females, selection should favor differential maternal investment in sons versus daughters depending on maternal condition and offspring reproductive potential. *Peromyscus maniculatus* and *P. polionotus* have a 25–28 day gestation period (Drickamer and Vestal 1973; Myers and Master 1983) that limits female reproductive potential; during this time, males have the opportunity to seek additional mating opportunities (Birdsall and Nash 1973; although *P. polionotus* rarely do [Foltz 1981]), which allows for greater reproductive variance in males than females, which meets one condition of the prediction. Our data, however, do not support the maternal resource allocation hypothesis because we found no association between testis size and maternal identity (*N*_*mothers*_=19; Figure S1), and no effect of litter order on testis size (*N*_*litters*_=140, 7-21 litters/female; Figure S2). In other words, we did not find that some females (e.g. females in greater condition) consistently produced sons with large testes, nor that they produced them earlier or later in life. Absence of a maternal effect in a captive laboratory setting is perhaps not surprising given that resources are fixed and food is provided *ad libitum*. Further evidence that maternal effects are unlikely here is that large litters in a congener, *P. californicus*, tend to be male-biased, potentially because males are slightly smaller at birth and less costly to produce than females (Cantoni et al. 2011), which is counter to the hypothesis that male quality and male-biased sex ratio may be both positively associated with maternal condition. Therefore, in our study, it is unlikely that the differences in testis size among males are due to maternal resource allocation and instead are due to differential investment in gamete production by the males themselves or passive exposure to increased testosterone *in utero*.

Our findings support the prediction that male testis size is phenotypically plastic and dependent on the early social environment. In placental mammals, the intrauterine position of embryos can influence the sexual development of individuals (Ryan and Vanderbergh 2002). In rats (van der Hoeven et al. 1992) and gerbils (Clark et al. 1993), male fetuses that develop between two brothers have larger testes at birth than males that are positioned between two sisters. The negative effects of intrauterine position on females associated with their reproductive potential (Ryan and Vanderbergh 2002), suggest that, at least in part, the intrauterine effect of brothers on testis size is likely due to passive uptake of circulating hormones *in utero*. However, it is also possible that males can actively respond to social cues of potential future competitors, either *in utero* or during the postnatal period prior to weaning, by increasing their investment in gamete production when reared with more brothers. For example, in the moth, *Plodia interpunctella,* larval male investment in reproductive development is associated with eventual mating frequency and risk of sperm competition (Gage 1995). In mammals, gamete production is more plastic in males compared to females, the latter of which are born with all the oocytes they will release during their lifetime. In male rodents, the first wave of spermatogenesis begins 4-7 days after birth (Russell et al. 1990), and differences in sperm production in adults is associated with sperm competition risk (Ramm and Stockley 2009; Firman et al. 2013; Klemme et al. 2014). Similar patterns in sperm production have been observed in *Peromyscus*: testis size is positively correlated with population density (Long and Montgomerie 2006) and length of breeding season (Ribble and Millar 1992). In this study, we found that testis size is not associated with the number of female siblings nor total litter size, suggesting that the critical component of population density that mediates investment in gamete production may be number of potential rivals. Similarly in humans, where intrauterine effects are reduced, men with more brothers produce significantly faster sperm (Mossman et al. 2013). Taken together, our results suggest that social conditions during early stages of development, either *in utero* or during the postpartum period, can have a critical influence on male reproductive potential.

In conclusion, we show that differential investment in sperm production may be both broadly shaped by selection over evolutionary timescales and also finely tuned by perceived competition risk during development. While our study does not allow us to directly test if greater testis size of males reared with more brothers is due to increased exposure to testosterone *in utero* or an active response to perceived competitors, the reproductive outcome is the same: males with more brothers develop larger testes, which is associated with greater sperm production and ultimately, fitness.

## DATA ACCESSIBLITY STATEMENT

Data are uploaded to Dryad.

## AUTHOR CONTRIBUTIONS

HSF and HEH designed the study. HSF, KAH and WDW collected data. KAH carried out the statistical analyses. All authors helped draft the manuscript and gave final approval for publication.

### ACKNOWLEDGEMENTS

We are grateful to Emily Jacobs-Palmer for her useful feedback on this study, and to Mollie Manier and Irene Liu, who provide helpful comments on this manuscript. This work is funding by a NICHD K99/R00 Pathway to Independence Award to HSF (R00HD071972), and a NSF Postdoctoral Research Fellowship to KAH (1711817). HEH is an Investigator of the Howard Hughes Medical Institute.

## Supporting information

**Supplementary Figure 1.**
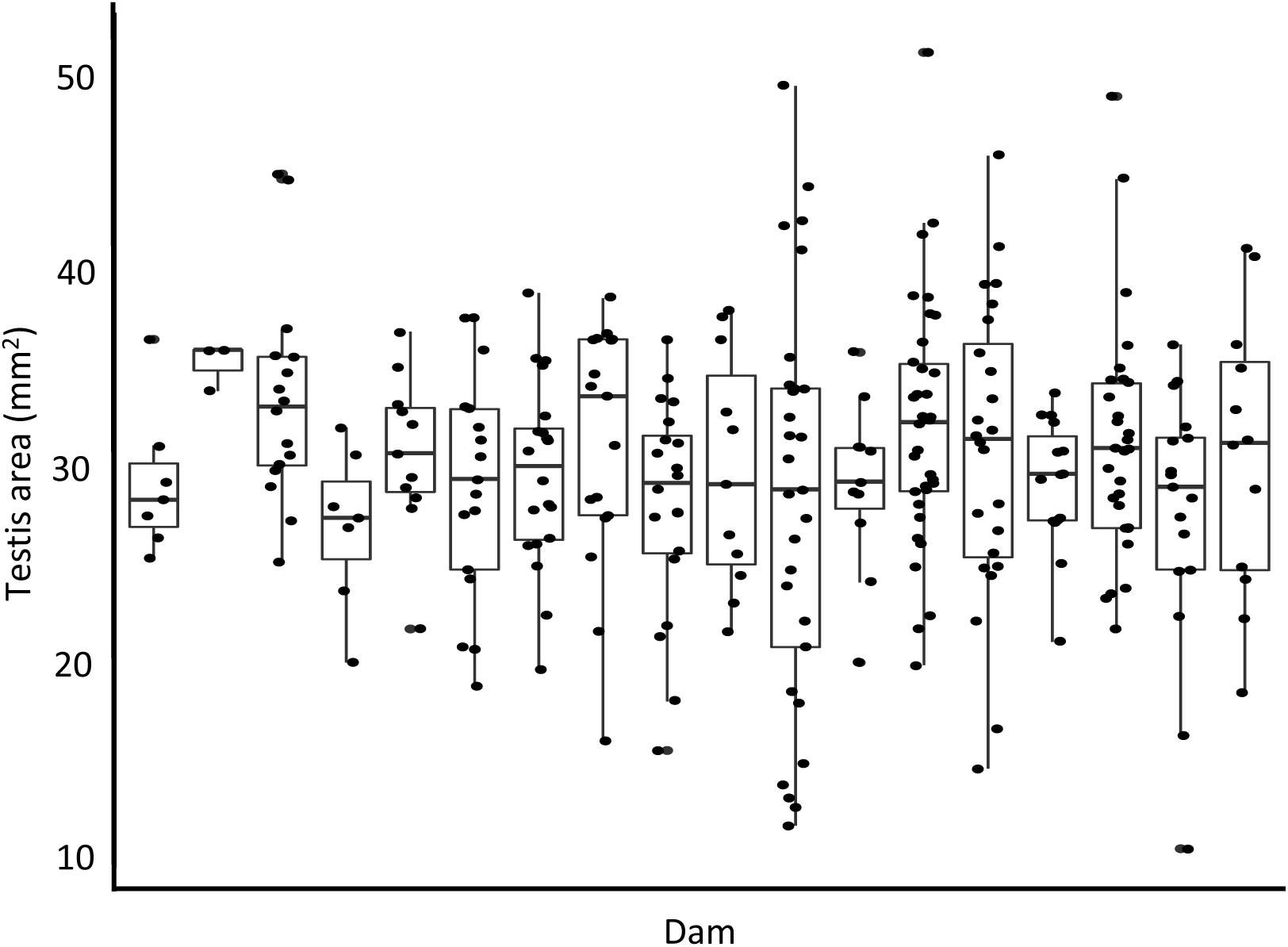
Mean testis size of F_2_ hybrid males sorted by maternal lineage (dam).

**Supplementary Figure 2.**
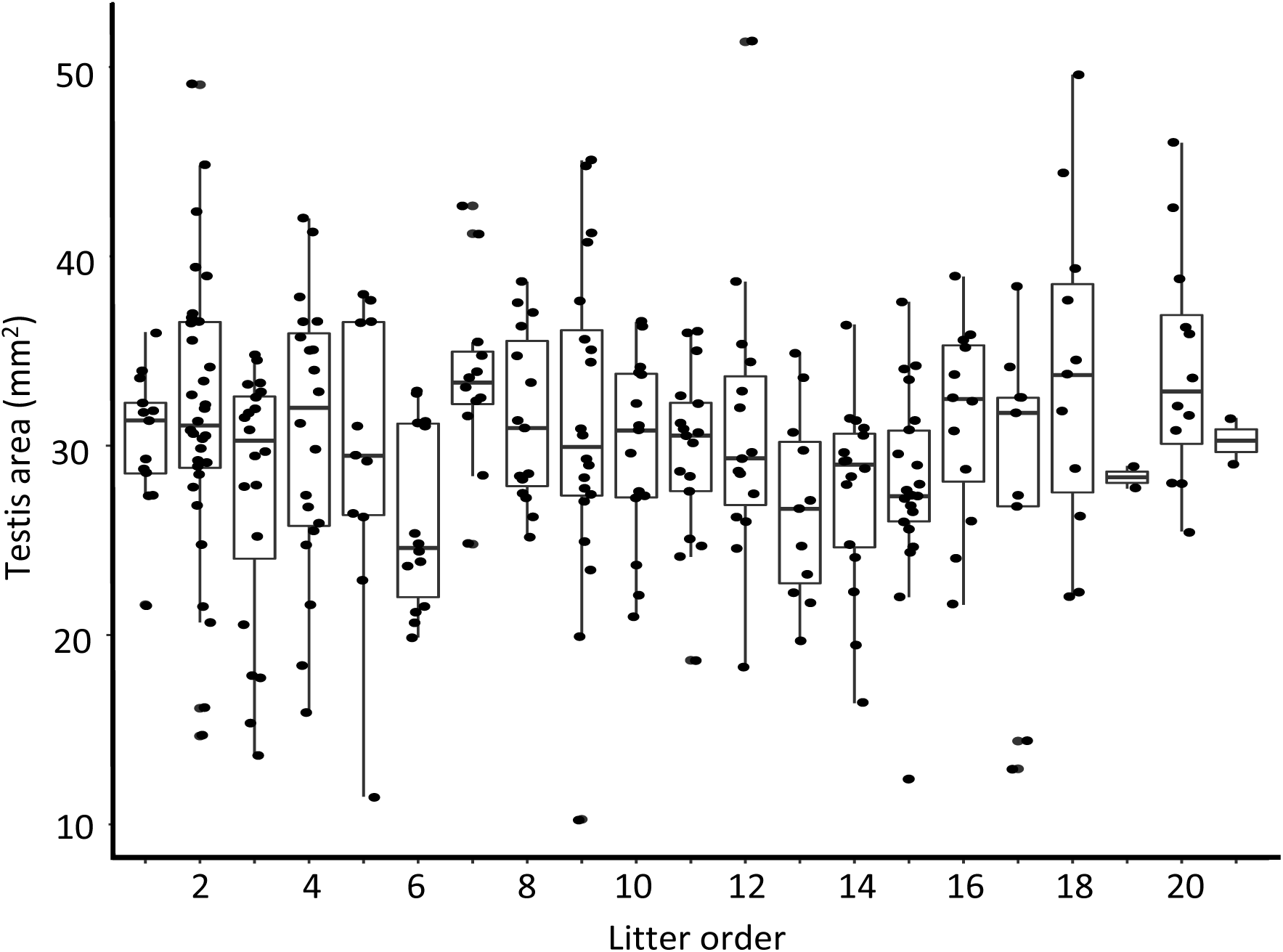
Mean testis size of F_2_ hybrid males sorted by litter order.

